# Integrative Multi-Omics Analysis Reveals Molecular Signatures of Age-Related Decline in Vervet Leg Skeletal Muscle

**DOI:** 10.64898/2026.09.25.754162

**Authors:** Maitree Patel, Eleanor V. Rudasi, Brett Frye, Jacob D. Negrey, Laura A. Cox, Ge Li, Matthew Jorgensen, Thomas C. Register, Kylie Kavanagh, Carol A. Shively, Ellen E. Quillen

## Abstract

Age-related decline in skeletal muscle function, often manifesting as slower gait speed, contributes significantly to frailty and loss of independence. To uncover the molecular basis of this decline, we investigated coordinated changes across the transcriptome, miRNome, and methylome in vastus lateralis muscle from female vervet monkeys (*Chlorocebus aethiops sabaeus*), contrasting middle-aged animals with normal gait speed versus older animals with declining gait speed. Through multi-omics integration, we identified a molecular signature that sharply distinguishes these two groups. This signature involves key regulatory molecules, including miR-181a-5p, miR-320b and miR-425 control of *SYNCRIP*, miR-193b-3p regulation of *MCL1* and *PLAU*, as well as a multi-omic regulatory network governing *ZNF274* and *ANP32E* gene expression, and altered levels of mRNAs such as *VEZT*. Functional annotation implicates miRNA signatures with inverse expression of their mRNA targets operating within systems of skeletal muscle biology, metabolism, and potential immune-inflammation mechanisms. Collectively, these findings reveal an interconnected network of epigenetic, post-transcriptional, and transcriptional alterations associated with impaired muscle function during aging. This molecular signature provides novel insights into potential mechanisms driving sarcopenia and highlights specific molecules as potential targets for interventions aimed at preserving muscle health.

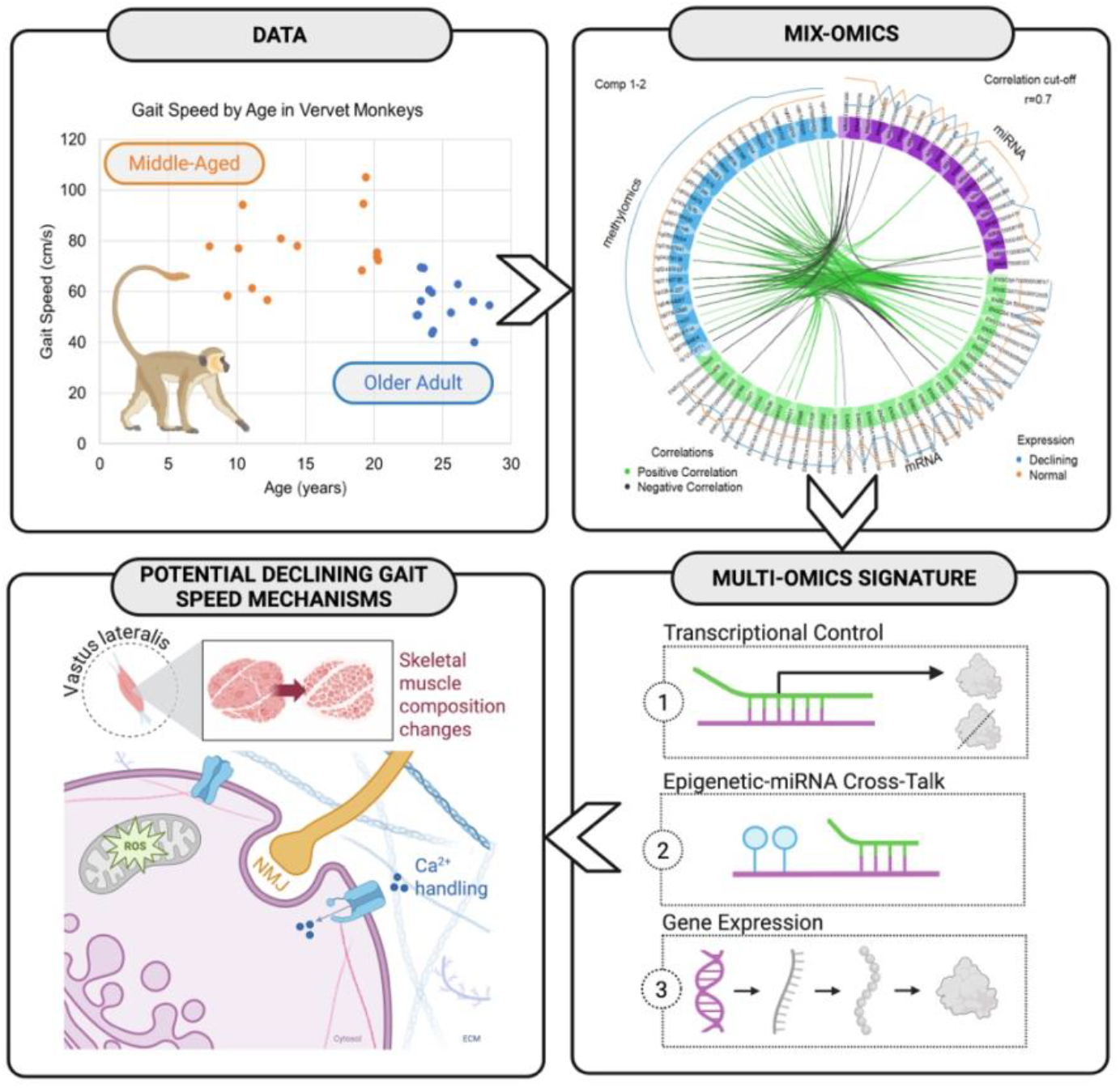

## Introduction

Gait speed has emerged as a robust biomarker of health and longevity in older adults, with slower walking speeds correlating strongly with higher morbidity [1, 2], reduced functional capacity [3], and early mortality [4, 5]. Human studies have linked gait speed to a range of age-related conditions, while evidence from non-human primates (NHPs) supports it as a translational biomarker of biological ageing. For instance, older female monkeys walk and climb stairs significantly more slowly than younger ones [6, 7], highlighting age-related decline in gait speed as a valuable phenotype for investigating the underlying mobility loss. While age and gait speed are highly correlated, focusing on the molecular changes associated with age may provide insight into the pathophysiology of impaired mobility.

Vervet monkeys (*Chlorocebus aethiops sabaeus*) offer a powerful model for investigating muscle aging. They share key physiological characteristics with humans, including similar genetic and gene expression profiles, comparable muscle structure, metabolic pathways, and aging trajectories, while controlled diet and housing conditions minimize confounding environmental variables [8]. The translational utility of vervets for muscle research is supported by studies demonstrating age-related reductions in radiographic muscle density [9], muscle cross-sectional area [10], and intrinsic force generation [11]. Age-related muscle atrophy and shifts in fiber-type composition in vervets also closely resemble human muscle aging [12], reinforcing their suitability for modeling sarcopenia and age-related mobility decline. In a sub-study of active vs sedentary old monkeys, sedentary monkeys had increased plasma LBP-1, suggestive of elevated microbial translocation independent of age, in non-sedentary monkeys’ insulin sensitivity was positively associated with frequencies of walking or running and negatively associated with LBP-1 [13].

Previous studies of gene expression changes in NHP skeletal muscle have identified pro-inflammatory transcriptomic changes with age [14], as well as proteomic and metabolomic alterations in protein turnover, energy metabolism pathways, and structure and function [15, 16]. Accumulating mitochondrial DNA mutations [17] and dysfunction are also central features of skeletal muscle aging in NHPs [18]. While these single-omics approaches have provided valuable insights, they offer only partial snapshots of the complex, interconnected biological processes driving muscle aging. Here we describe a multi-omic approach to compare skeletal muscle between middle-aged animals with normal gait speed and older animals with declining gait speed. By analyzing transcriptomic, small RNA, and methylation changes from the same skeletal muscle samples drawn from well-characterized female vervet monkeys, this integrated strategy allows us to decipher the complex molecular signatures of age-related functional decline in muscle.

## Methods

### Study Subjects and Gait Speed Assessment

This study analyzed 28 adult female vervet monkeys (*Chlorocebus aethiops sabaeus*), aged 8 to 29 years (maximal lifespan 31 years [19]), selected from the Vervet Research Colony (VRC) at Wake Forest University School of Medicine. The VRC is an NIH-supported breeding colony housing animals under controlled conditions, extensively characterized for aging, cardiometabolic, and neuroscience research. Animal housing and husbandry practices adhered to the guidelines of the Animal Welfare Act and the Guide for the Care and Use of Laboratory Animals. All procedures were approved by the Wake Forest Institutional Animal Care and Use Committee (A24-149) and complied with all relevant United States and North Carolina regulations.

Physical performance was assessed using gait speed, measured as previously published [6]. Briefly, each monkey’s typical, unprompted walking pace was timed over five independent trials, and the average speed (cm/sec) was calculated. Based on the gait speed measurements, animals were retrospectively categorized into two equal groups (n=14 per group): “Normal” gait speed, reflective of the common gait speed across animals in young adulthood, and “Declining” gait speed, seen among older animals, for the purpose of supervised multi-omics analysis (Fig1). While there is a significant difference in mean gait speed between these two groups (21.7 cm/sec, *p* = 4.5x10^−5^), there is overlap in gait speed between the groups and a correlation between age and gait speed (R^2^ = 0.25). Therefore, we cannot completely disentangle age and gait in this analysis but should consider the comparison to be between younger animals and older animals with a cutoff age of 21, approximately when gait speed begins to decline.

**Fig 1.**
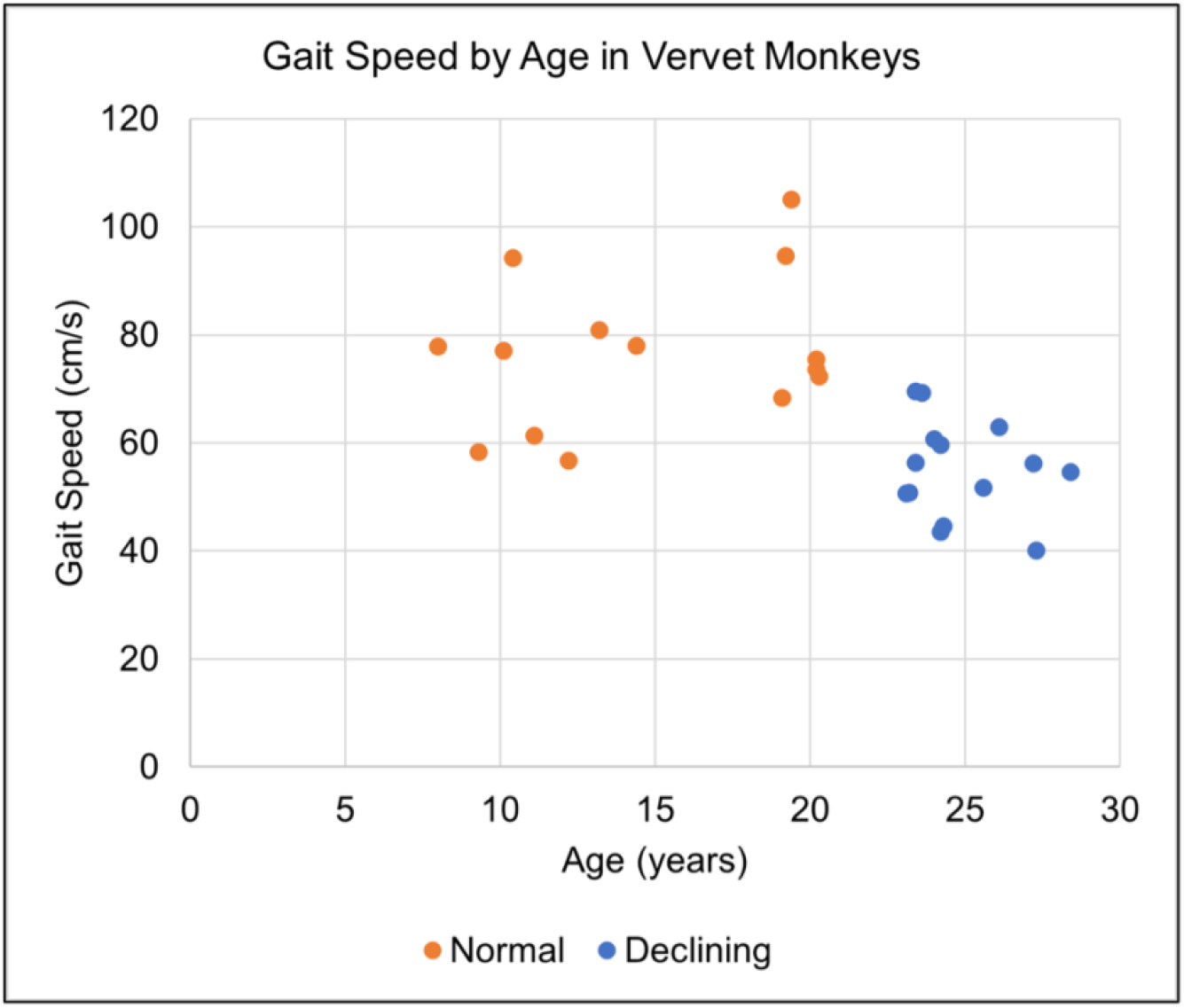
Gait Speed by Age in Vervet Monkeys Scatter plot illustrating the relationship between age (years) and gait speed (cm/sec), with orange points indicating those with “Normal” gait speed and blue points indicating those classified as “Declining.”

### Tissue Collection and Omics Data Generation

*Vastus lateralis* biopsies were collected while the vervet monkeys were sedated for routine health checks. Samples were snap-frozen and stored at -80°C until extraction of total RNA and genomic DNA from approximately 5mg of homogenized, frozen muscle using Qiazol Lysis Reagent (Illumina) according to the manufacturer’s instructions. DNA and RNA integrity and concentration were determined by an Agilent 2100 Bioanalyzer and UV-Vis Spectrophotometer (Nanodrop 2000). DNA methylation was assessed using the Illumina Infinium MethylationEPIC BeadChip array of 866,895 CpG sites using standard protocols run by the commercial lab Diagenode. Raw methylation data from the EPIC array was processed using the ChAMP package [20]. Default filters for non-CG probes, SNPs, XY chromosomes, and multi-hit probes were disabled during loading and filtering was performed based on a vervet-specific annotation [21]. Default quality control assessments, probe normalization, and batch correction were performed. No technical batch effects were detected. Raw and processed data files along with metadata for all methylation and mRNA/miRNA data generated during this study are currently in the process of being submitted to NCBI Gene Expression Omnibus (GEO) and Sequence Read Archive (SRA).

cDNA libraries were prepared for combined mRNA and small RNA sequencing using the NEXTFLEX Combo-Seq mRNA/miRNA kit (Perkin Elmer). RNA sequencing was performed on the Illumina NovaSeq 6000 instrument using reagents provided by the manufacturer. All trimming, quality control, alignment, and quantification were performed in Partek Flow (Qiagen). mRNAs were aligned to the vervet genome (Chlorocebus_sabeus 1.1) using the STAR aligner [22] with a Phred quality score ≥ 30. smRNAs were aligned to the human miRBase database [23] using Bowtie 2 [24]. Following standard post-alignment quality control metrics, mature miRNA, transcript, and gene-level counts were generated. Genes with low counts across samples were filtered and remaining counts were normalized using the Trimmed Mean of M-values (TMM) method and converted to log2 counts per million in edgeR [25]. miRNA counts were similarly normalized.

### Cell Type Deconvolution using CIBERSORTx

To account for potential confounding effects of cellular heterogeneity, the relative proportions of major cell types were estimated using the CIBERSORTx deconvolution algorithm [26] in reference to a validated signature matrix derived from human skeletal muscle single-cell/nucleus RNA-seq data [27]. CIBERSORTx output provides relative fractions of eight cell types (endothelial cells, fibroblasts, lymphoid cells, myeloid cells, myocytes, pericytes, satellite cells, smooth muscle cells) for each sample. Differences in estimated cell proportions between the Normal and Declining gait speed groups were assessed using independent two-sample t-tests. Cell types showing significant differences (p < 0.05) were included as covariates in downstream differential analyses of mRNA and DNA methylation data.

### Integrative Multi-Omics Analysis using DIABLO

Integration of the datasets was performed using the DIABLO framework within the mixOmics R package [28, 29]. Feature lists were filtered to the top 9,999 differentially methylated CpG sites that had a corresponding human gene annotation and top 9,999 differentially expressed mRNAs. Only 194 variable miRNAs were annotated using the human miRBase, so all were included in the analysis. Prior to building the final model, pairwise relationships between the omics datasets were explored using sparse Partial Least Squares (sPLS) analysis. A design matrix was specified where all off-diagonal elements were set to 0.1, indicating an expectation of moderate correlation between all pairs of omics layers. To optimize classification performance (by minimizing balanced error rate [BER]), the number of components and features per component for each dataset were determined using 10-fold cross-validation with 50 repeats. This tuning indicated an optimal model included 2 components. The final model’s performance was assessed for overall error rate (ER), BER, and area under the ROC curve (AUC). Diagnostic plots including sample projections, feature loading weights, a clustered image map, a circos plot, and a relevance network were generated to evaluate model fit, component structure, and feature relationships.

### Prediction of Molecular Interactions

Putative mRNA targets for the miRNAs selected by the final DIABLO model were predicted using the miRNAtap package [30]. Predictions were aggregated across TargetScan, miRanda, PicTar, DIANA, and miRDB databases using the geometric mean method, requiring target prediction from at least two sources. As miRNAtap uses human annotations, predicted target EntrezGene IDs were converted to human gene names using the biomaRt package (v2.60.1) [31, 32]. DIABLO-selected vervet mRNA transcript IDs were mapped to vervet gene names using biomaRt (csabaeus_gene_ensembl dataset, ChlSab1.1). Overlaps between the DIABLO-selected vervet gene list and the human predicted miRNA target gene list were identified based on shared gene names.

Potential cis-regulation of mRNAs and miRNAs by methylation was assessed by CpG site annotations relative to gene features and CpG islands in proximity to the transcripts of interest. Potential indirect interactions of joint effects of methylation and miRNAs were explored by comparing annotation of DIABLO-selected CpGs with predicted targets of DIABLO-selected miRNAs from the miRNAtap/biomaRt analysis.

### Functional Enrichment of DIABLO-Selected Multi-omics Molecular Features

To investigate the biological functions underlying interconnected multi-omics signatures, Gene Ontology (GO) and Kyoto Encyclopedia of Genes and Genomes (KEGG) enrichment analysis was performed in R. Target genes for each miRNA was predicted using miRNAtap (v1.40.0). miRNA-mRNA pairings with concordant expression levels in our cohort that had experimental evidence from the Human TargetScan source database [33] were summarized using QIAGEN IPA (QIAGEN Inc., https://digitalinsights.qiagen.com/IPA). Entrez gene lists corresponding to each DIABLO selected omics were analyzed using the clusterProfiler (v4.14.6) [34] package in R with “hsa” as the organism code. The pathway results were interpreted based on the overall biological relevance given the phenotype under investigation, with an adjusted p-value < 0.05 and with >= 2 genes matched within it.

## Results

### Estimated Muscle Cell Type Composition Differs Between Age-Associated Gait Speed Groups

Cellular heterogeneity using CIBERSORTx yielded strong goodness-of-fit between the observed and predicted cell-type proportions (Pearson correlations 0.70–0.93, p-value < 0.05). T-tests comparing the estimated proportions between gait speed groups revealed significantly lower proportions of endothelial cells (p ∼ 0.034) and significantly higher proportions of myocytes (p ∼ 0.013) in the Declining group compared to the Normal group (Fig2). Full estimated cell type proportions are available in the Supplementary Materials.

**Fig 2.**
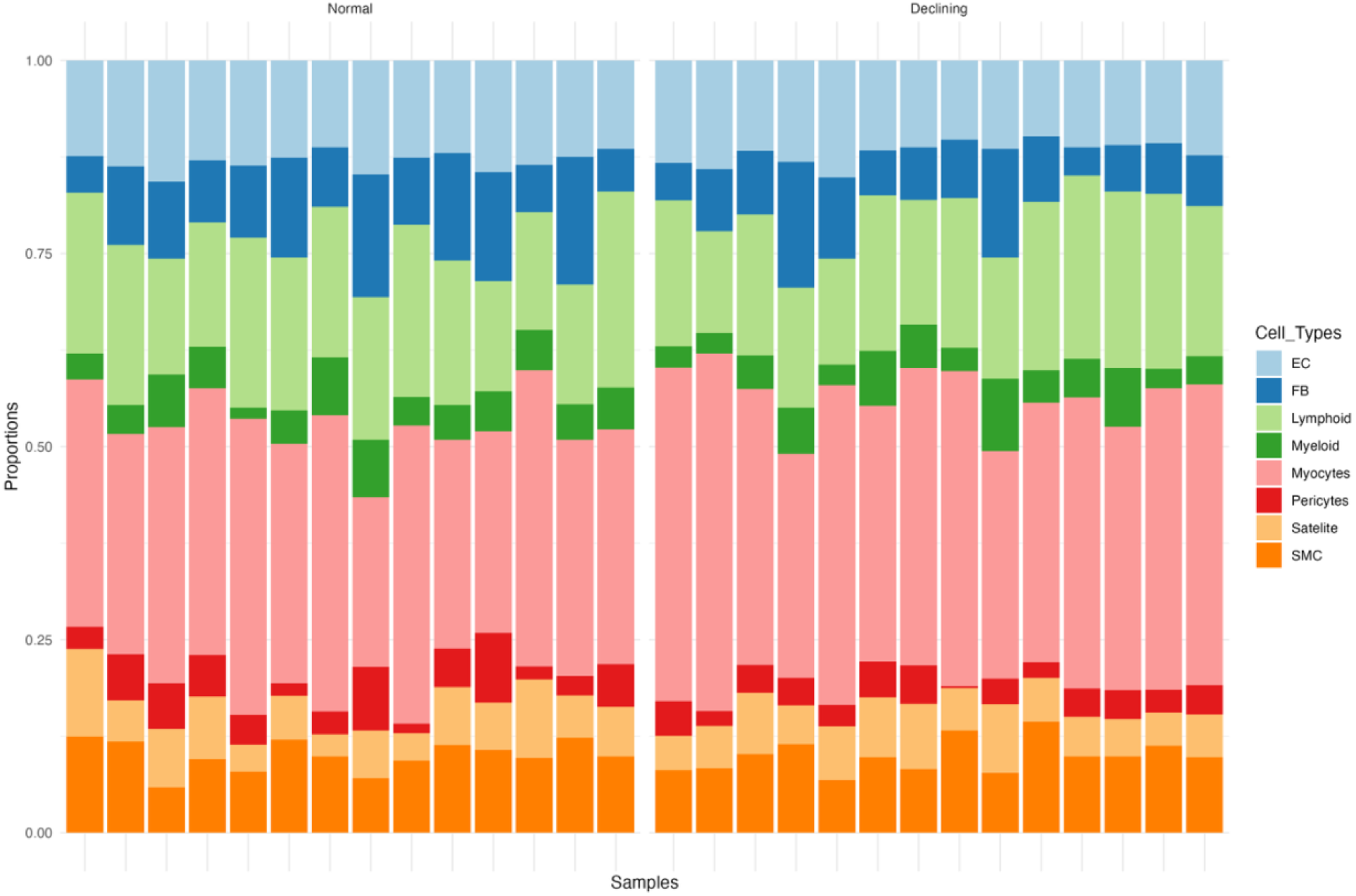
Estimated Cell Type Proportions in Vervet Vastus Lateralis Muscle Biopsies via CIBERSORTx Deconvolution Stacked bar chart displaying the estimated relative cell type composition for each individual female vervet’s Vastus Lateralis (VL) muscle biopsy sample (n=28). Each bar represents one animal, with segments showing the relative percentage (y-axis) of different cell types, summing to 100%. The legend indicates the cell types represented by each colored segment (EC: Endothelial Cells, FB: Fibroblasts, SMC: Smooth Muscle Cells).

### Single-Omics Analyses Identify Limited Features Associated with Gait Speed Classification

To assess molecular differences using traditional single-omics approaches, DE analysis of mRNAs and miRNAs, and differential methylation analysis of CpG sites were performed between the Normal and Declining groups while adjusting for cell type heterogeneity. DE analyses revealed only a few significant features. We found a single differentially expressed gene between the gait speed groups, the *MYL10* gene (FDR < 0.05, absolute LogFC ∼ -4). miRNA DE analysis revealed only two significant features, including miR-100-5p and miR-223-3p, both having LogFC < 2. All-in-all, the single omics analysis revealed only limited biological signals underlying gait speed decline in female vervets. And lastly, three CpG sites were hypermethylated in the declining gait speed group with a mean difference in beta ∼ 1, they include cg20410171 (near *ZNF578*) cg11833660 (near *SSH1*), and cg00077017 (near *FLNB*). Comprehensive descriptions of significant single-omics features can be found in Supplementary Materials.

### Integrative Multi-Omics Analysis using DIABLO Reveals a Robust Signature Discriminating Gait Speed Groups

The DIABLO framework integrated the pre-filtered mRNA (9,999 features), miRNA (194 features), and DNA methylation (9,999 features) datasets to identify a multivariate signature that discriminated between the Normal and Declining gait speed groups. Initial pairwise sPLS analyses confirmed strong correlations between the datasets (Figa) and the final DIABLO model, tuned via cross-validation to include two components, demonstrated high classification accuracy. The overall Balanced Error Rate (BER) was low (∼17 – 18% across components, centroids distance), and the Area Under the ROC Curve (AUC) based on the miRNA block, very high (AUC = 0.9898), indicating strong predictive ability. Diagnostic plots confirmed good separation between the Normal and Declining groups based on sample projections onto the model components and strong concordance between the latent structures derived from the different omics blocks, particularly for Component 1 (inter-block variate correlations R = 0.74–0.90, Fig3a). These correlations (R = 0.90 for miRNA vs mRNA; R = 0.74 for miRNA vs methylomics; R = 0.86 for mRNA vs methylomics) demonstrate the strong concordance achieved between the latent structures derived from the different omics datasets in the final integrated model. A heatmap of the selected features showed clear clustering of samples according to gait status (Fig3b). Several DIABLO-selected miRNAs are predicted to target DIABLO-selected mRNAs based on miRNAtap predictions, which suggests that these components are capturing functionally integrated pathways. (**Error! Reference source not found**.). Particularly, three miRNAs (miR-181a-5p, miR-320b, and miR-425) targeted the same gene, Synaptotagmin Binding Cytoplasmic RNA Interacting Protein (*SYNCRIP*). miR-181a-5p (MIMAT0002506) and miR-425 (MIMAT0006324) contributed a reliable amount on component 1 (Supplementary Fig3a) and miR-320b (MIMAT0024312) on component 2 in discriminating between gait speed groups (Supplementary Fig3b). Direct comparison of gene names associated with DIABLO-selected CpG sites and DIABLO-selected mRNA transcripts yielded no overlaps. Surprisingly, integrating DIABLO-selected CpG sites (n=21) and DIABLO-selected miRNAs (n=20) through their shared target/associated genes identified potential indirect regulatory links (**Error! Reference source not found**.). Four genes (*KCNH6, BCL11B, ZNF274, ANP32E*) nearby CpGs in the two DIABLO components are also predicted targets of selected miRNAs. Both *ZNF274* and *ANP32E* are associated with age and reveal key molecular mechanisms underlying functional decline in skeletal muscles.

**Fig 3.**
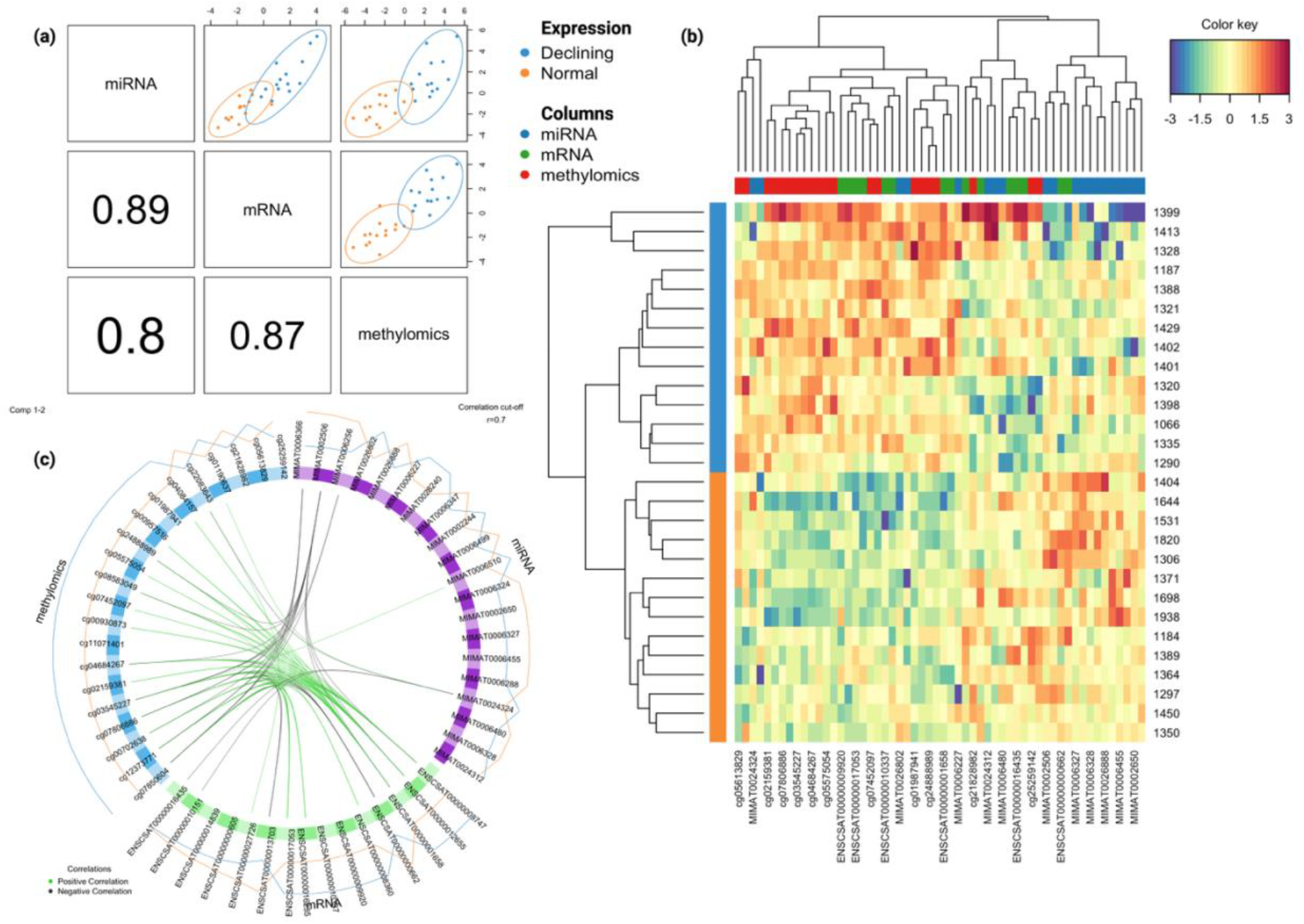
DIABLO Model Results (a) Diagnostic plot visualizing the final DIABLO model integrating mRNA, miRNA, and DNA methylation data. Upper right triangle panels: Scatter plots show individual samples projected onto the first two components derived by the model, viewed within each omics block. Samples are colored by gait status group (Declining = blue, Normal = orange), with 95% confidence ellipses indicating group separation achieved based on the integrated components. Lower triangle panels: Pearson correlation coefficients (R values) between the sample scores (variates) on the first component for each pair of omics blocks. (b) Heatmap displaying the relative abundance levels of the top features selected across mRNA, miRNA, and DNA methylation datasets by the final DIABLO model for all 28 vervet samples. Rows represent individual animals (labeled by ID) and are clustered without supervision based on their overall multi-omics profile similarity (dendrogram on left). The adjacent sidebar indicates the predefined gait status group for each sample. Columns represent the selected features (labeled by ID) and are clustered by profile similarity across samples (dendrogram on top). The sidebar beneath the column dendrogram indicates the omics type of each feature (Columns legend: Blue=miRNA, Green=mRNA, Red=methylomics). Heatmap color intensity corresponds to the scaled and centered abundance level for each feature in each sample. (c) The circos plot displays correlations exceeding 0.7 between variables from methylomics in teal, miRNA in purple, and mRNA in green. Each segment within a quadrant represents a specific feature variable (e.g., a CpG site, miRNA, or mRNA transcript). Green connection links represent positive correlations, while black links represent negative correlations. The outer lines reflect the expression level associated with gait status: blue for “Declining” and orange for “Normal.” The model selected features across all three omics types that contributed to group discrimination (Component 1: 10 mRNA, 16 miRNA, 16 CpG; Component 2: 5 mRNA, 5 miRNA, 5 CpG). Annotations and loadings of each feature on the two components are in Supplementary Materials. Visualization of the relationships between these top features via Circos plot (Fig3c) and relevance network (Supplementary Materials) revealed a highly interconnected signature spanning the different molecular layers that was not detected in the single omics analysis.

### Enrichment Analyses Identify Skeletal Muscle, Endocrine, and Immune Systems

KEGG and GO functional enrichment analysis revealed that miRNA-mRNA pairings with opposing expression signals enriched within significant pathways (p < 0.05, Count >= 2) of the skeletal muscle composition, immune and endocrine system, and metabolism. Effect of aging on skeletal muscle structure and dynamics are evident through 20 unique GO enrichment results surrounding muscle cell biology including proliferation, growth, migration, and development; most notable of them include regulation of skeletal muscle cell differentiation (G:2001014), skeletal muscle organ development (GO:0060538) and tissue development (GO:0007519), skeletal muscle tissue development (GO:0007519), and striated muscle cell migration (GO:0014909) and differentiation (GO:0051146). Immune system KEGG pathways include chemokine signaling (hsa04062), leukocyte transendothelial migration (hsa04670), Th17cell differentiation (hsa04659), and T cell receptor signaling (hsa04660) to name a few, together with four distinct GO enriched terms suggests a potential inflammation signal associated with aging skeletal muscles. Gene enrichment in systems of insulin action and signaling in addition to other metabolic and endocrine system (potential hormone imbalance) pathways further highlight the potential downstream effects of declining skeletal muscle dynamics and inflammation. The cholinergic synapse (hsa04725) KEGG enrichment result supports a neuromuscular junction (NMJ) link in aging vervets.

## Discussion

Declining gait speed is a clinically significant indicator of reduced functional capacity and increased health risks associated with aging. This study employed an integrative multi-omics approach to uncover the molecular underpinnings of age-related gait speed decline in the skeletal muscle of female vervet monkeys, a relevant non-human primate (NHP) model. By combining mRNA, miRNA, and DNA methylation data using the DIABLO framework, we successfully identified a robust multivariate signature capable of discriminating between middle-aged animals with normal gait speed and older animals with declining gait speed, achieving high classification accuracy (BER ∼16-19%, AUC >0.97). This success contrasts with traditional single omics DE analyses which identified a few significantly altered mRNAs, miRNAs, and CpG sites following correction for multiple testing (Online Resource Individual Omic Feature Analyses Results section). This highlights a key strength of our integrative strategy: its ability to capture subtle, coordinated changes across multiple molecular layers that collectively drive the phenotype, even when individual feature alterations do not reach statistical significance in isolation.

The multi-omics signature identified by DIABLO (Table S3-5; ESM4-ESM6) points towards specific molecular mechanisms potentially driving functional decline. Central to this signature is the gene *SYNCRIP* targeted by three DIABLO selected miRNAs: miR-181a-5p, miR-320b, and miR-425; DIABLO-selected mRNA *VEZT* (selected on component 1 and a target of miR-15a-5p), and CpG-miRNA (Table 2) regulatory interactions on genes *ZNF274* and *ANP32E*. Furthermore, miR-193b-3p (MIMAT0006227) which was a major contributor on component 1 targeted two genes, *MCL1* and *PLAU*. Most of these interactions have not been elucidated in sarcopenia models and therefore act as key potential novel biomarkers for predicting functional skeletal muscle decline in aging sarcopenia models.

**Table 1.**
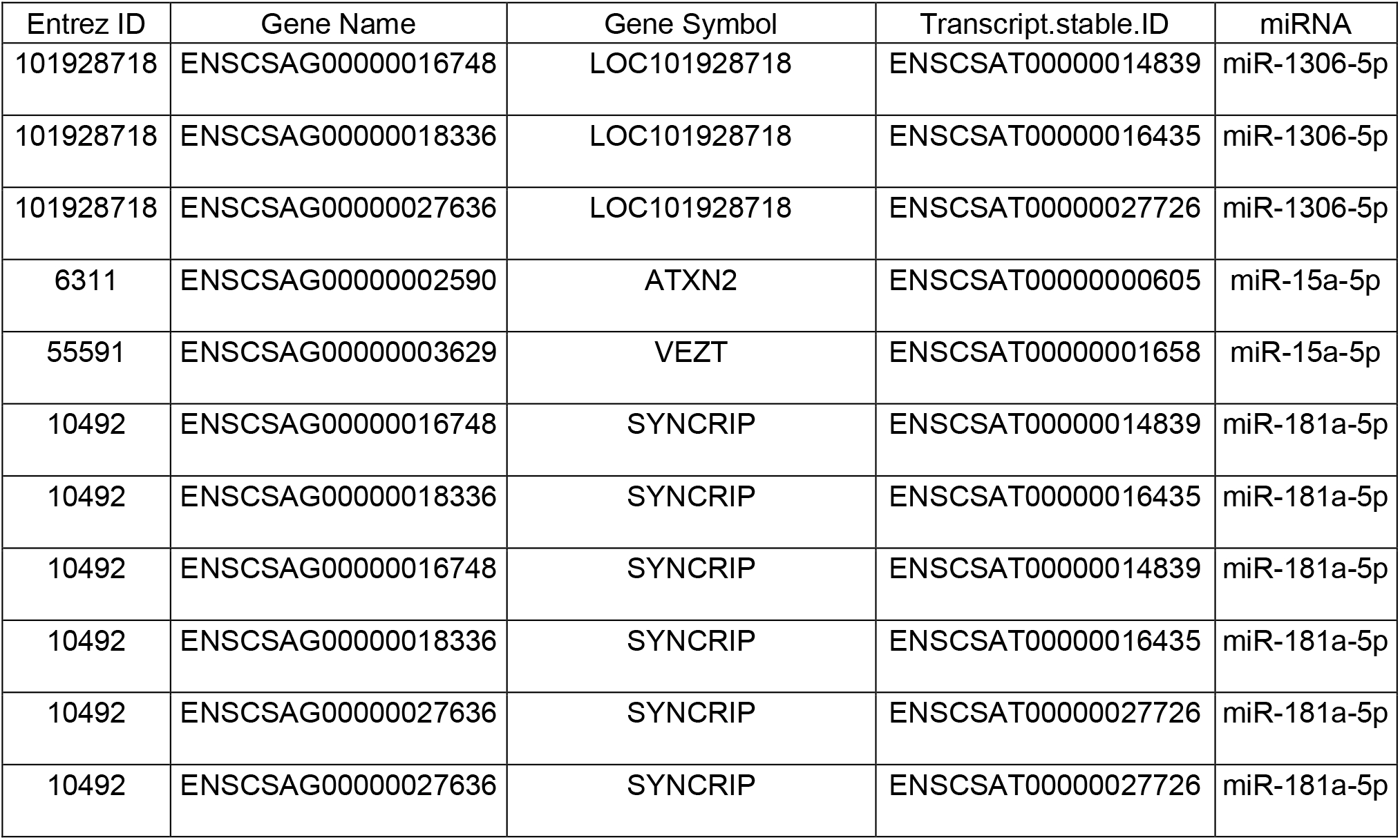
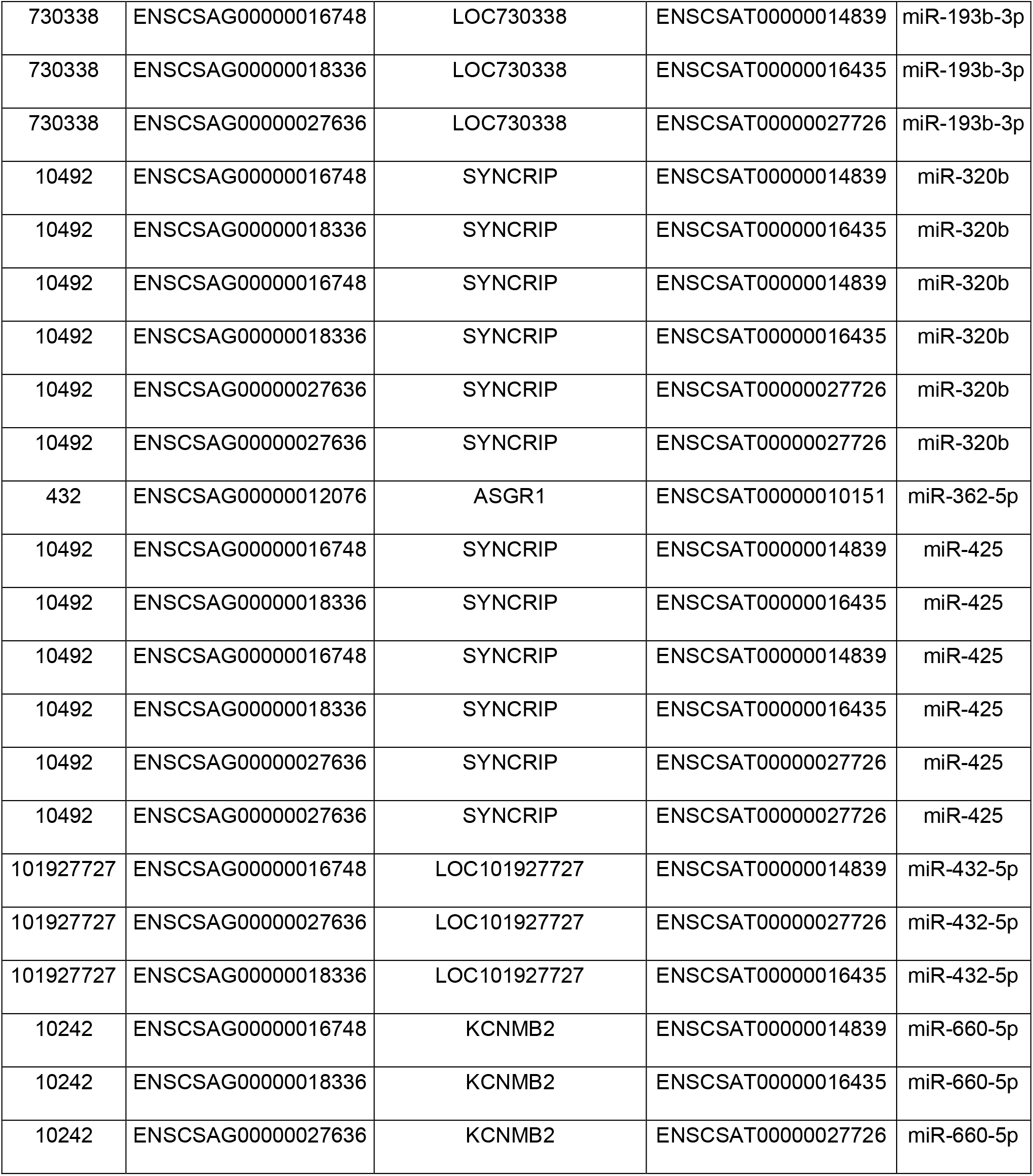
Predicted Interactions Between DIABLO-Selected mRNAs and miRNAs. The predicted interactions between mRNA transcripts selected by the DIABLO model (components 1 and/or 2) and miRNAs potentially regulating them. Putative mRNA targets of miRNAs were identified using miRNAtap (based on human annotations, requiring ≥2 database sources), and overlaps were found with DIABLO-selected vervet mRNAs mapped to gene names via biomaRt. The table lists the target mRNA EntrezGene ID, Gene Name, Ensembl IDs, gene description, and the corresponding predicted interacting miRNA. These interactions highlight potential post-transcriptional regulatory pathways involving key components of the identified multi-omics signature associated with gait speed status. Note that SYNCRIP is predicted to be targeted by miR-181a-5p, miR-320b and miR-425.

**Table 2.** Predicted CpG-miRNA Interactions via Shared Associated Genes Potential regulatory interactions involving CpG sites and miRNAs selected by or relevant to the DIABLO model. Interactions were identified by finding genes (Gene Name, Entrez ID) annotated to DIABLO-selected CpG sites (CpG Name) that were also predicted targets of specific miRNAs (miRNA, identified by name and MIMAT ID Alias), based on miRNAtap predictions. This analysis highlights instances where epigenetic (DNA methylation) and post-transcriptional (miRNA) regulation may converge on the same gene targets within the multi-omics signature associated with gait speed status.

| CpG Name | Gene Name | Entrez ID | miRNA Name | Alias |
| --- | --- | --- | --- | --- |
| cg01190637 | KCNH6 | 81033 | mml-miR-432-5p | MIMAT0006327 |
| cg01987941 | BCL11B | 64919 | mml-miR-15a-5p | MIMAT0002650 |
| cg04684267 | ZNF274 | 10782 | mml-miR-181a-5p | MIMAT0002506 |
| cg21828982 | ANP32E | 81611 | mml-miR-487b-3p | MIMAT0006347 |
| cg21828982 | ANP32E | 81611 | mml-miR-502-3p | MIMAT0006366 |
| cg21828982 | ANP32E | 81611 | mml-miR-767-5p | MIMAT0006510 |

The *SYNCRIP* gene encodes a nuclear ribonucleoprotein that regulates mRNA metabolism and might be associated with multiple muscle and neuromuscular disorders [35, 36] manifested as muscle weakness through direct functional declines in the muscle or indirectly through the dysregulation of nerve-muscle communication. This is further strengthened by abnormal gait and other movement related declining characteristics in individuals with SYNCRIP-related neurodevelopmental disorder (SYNCRIP-NDD) [37] in addition to other age-related clinical symptoms. The convergence of three DIABLO-selected miRNAs onto *SYNCRIP* points to a key putative regulatory axis driving gait speed decline. Furthermore, miR-181a-5p has already been associated with gait speed changes in clinical contexts [12], suggesting its regulatory role in physical performance within muscle tissues.

The *VEZT* gene encodes transmembrane protein vezatin that localizes in the adherens junction and is involved in its maintenance in adult life [38]. Though it is most associated and widely studied in relation with gastric cancer [39] and embryonic death [40], it has been associated with decrease locomotor activity through four phenotypes in mouse models by the International Mouse Phenotyping Consortium [41] and is a part of the pathway miR-targeted genes in the muscle cell [42]. Being the second largest contributing feature (ENSCSAT00000001658) on component 1 (> 0.5 loadings weight), *VEZT* expression signal might suggest reduced muscle function through shifts in the adherens junction biology at the NMJ.

*MCL1* and *PLAU* are both gene targets of miR-193b-3p, with confident IPA pairing predictions with mRNA targets of concordant expression values. Widden and Placzek [43] highlight the non-apoptotic functions of *MCL1*, including cell proliferation, calcium handling, and mitochondrial dynamics [44]. Calcium ion transport controls muscle contractions [45] and is implicated in skeletal muscle aging [46]. A key observation was the already established associations of MCL1 with multiple white blood cell (WBC) traits including basophil [47, 48] and monocyte counts [49, 50], neutrophil to lymphocyte ratio [51, 52] among others which might be indicative of blood contamination of samples or simply an overall aging signature. *PLAU* has also been identified as one of the 16 senescence associated secretory phenotypes in smooth muscle cells and muscle stem cells by Li, Li [53] to particularly identify therapeutic targets for sarcopenia. miR-149b-3p regulation of both these genes might trigger the onset of skeletal muscle related aging mechanisms including mitochondrial dysfunction and cellular senescence.

We also identify *ZNF274* and *ANP32E* as potential regulatory interactions involving DIABLO selected CpGs and miRNAs. Both have been linked to aging-related functional differences [54-56]. *ANP32E* is additionally involved in muscle regeneration [57] and is a molecular chaperone regulating proteostasis and chromatin remodeling during age-related muscle decline in *YTHDF2*-deficient mice [58]. It is important to note that these features might be suggestive of molecular systems underlying aging in general, since it is highly correlated with our phenotype. Additionally, *ANP32E* association with lymphocyte percentage [52] might also reflect blood contamination of samples. Although these molecular players have well-established biological functions, and some are implicated in aging, their interconnected role in sarcopenia remains largely unexplored, highlighting the need for experimental validation.

Pathway analysis of DIABLO-selected miRNA targets of genes with concordant expression levels in our data reveals a holistic mechanistic overview of skeletal muscle aging underlying gait speed decline (Table S1 and S2; ESM2-ESM3). The overwhelming GO enrichment signals surrounding muscle biology might suggest physical changes in fiber type and size [59] in our cohort and the consequent reduction in muscle function. The latter is also supported by the enrichment of genes within various significant endocrine pathways. Since muscle is an organ that regulates and facilitates homeostasis; a reduction in its function can point towards endocrine dysfunction. Furthermore, the enrichment of immune system pathways might underscore a systemic inflammation signature accompanying aging skeletal muscles. Other significant pathway enrichment results are also reflective of skeletal muscle aging in sarcopenia, namely, cellular senescence (hsa04218), longevity regulating pathway (hsa04213), insulin signaling pathway (hsa04910), adherens junction (hsa04520) and potentially the AGE-RAGE signaling pathway in diabetic complications (hsa04933). Taken together, our pathway enrichment results show that muscle composition changes, glucose metabolism, inflammation, and cellular stress are intrinsic to functional decline in aging skeletal muscles and might underly the multi-omics signature we identified.

Our analysis also accounted for cellular heterogeneity by estimating cell type proportions using CIBERSORTx. The finding that the Declining group had significantly lower estimated endothelial cell proportions and higher myocyte proportions compared to the Normal group is noteworthy. While adjusted for in subsequent analyses, this difference itself may reflect underlying biological changes associated with sarcopenia, such as potential alterations in muscle capillarization or fiber type composition [60, 61], although further investigation with specific histological or single-cell methods would be needed to confirm this.

It is important to consider the phenotype definition in this study. We categorized animals based on gait speed, a functional outcome strongly associated with age in this colony [6] and predictive of morbidity and mortality in humans [4, 5]. While our analysis successfully identifies molecular signatures linked to this functional decline, the strong correlation between age and gait speed makes it difficult to completely disentangle effects purely driven by ‘slowness’ versus those driven by the broader biological aging process in this observational design. Therefore, the identified signature is best interpreted as representing the molecular landscape associated with age-related functional decline in vervet skeletal muscle.

This study has several strengths, including the use of a well-characterized NHP model relevant to human aging, the application of a powerful multi-omics integration technique (DIABLO), and the adjustment for cellular heterogeneity. However, limitations must be acknowledged. The sample size (n=28) is modest for multi-omics studies, although typical for NHP research. The study included only female animals, potentially limiting generalizability. Our reliance on a vastus lateralis muscle biopsy may not fully represent systemic muscle aging. Furthermore, the cell type deconvolution and miRNA target/GO analyses relied on human annotations and signature matrices due to the lack of vervet-specific resources, introducing potential cross-species inaccuracies that warrant caution in interpretation. Finally, this is an observational study identifying correlations; experimental validation is required to establish causal links between the identified biomarkers and functional decline.

Future research should focus on validating the key novel biomarkers (e.g., miRNA regulation of *SYNCRIP* gene, miR-193b-3p targets *MCL1* and *PLAU*, gene expression, *ZNF274- and ANP32E*-associated CpGs and miRNA targets) and predicted interactions using targeted molecular techniques and functional assays in cell or animal models. Longitudinal studies tracking these molecular changes alongside functional decline within individual animals would strengthen causal inference. Expanding the cohort to include male animals and potentially other muscle types would enhance generalizability. Incorporating additional omics layers, such as single-cell multi-omics approaches and broader proteomics, could provide crucial information on cellular phenotypes and transitions as well as protein abundance and post-translational modifications downstream of the identified transcriptomic and epigenetic signatures. Advanced mass spectrometry-based platforms, such as the Seer platform, offer the potential for deep proteome coverage and the ability to precisely map protein-altering variants, complementing the findings of this study [62, 63].

In conclusion, this study successfully employed an integrative multi-omics strategy (DIABLO) to identify a robust molecular signature associated with age-related gait speed decline in vervet monkey skeletal muscle, even when traditional single-omics analyses yielded insufficient and independent biological features after multiple testing correction. The identified signature involves interconnected changes across mRNA, miRNA, and DNA methylation levels, highlighting key biomarkers such as miR-181a-5p, miR-320b, and miR-425 regulation of *SYNCRIP*, and specific CpG sites and miRNAs potentially regulating *ZNF274, ANP32E; and* miR-193b-3p control *of MCL1 and PLAU*, linked through complex regulatory networks. These findings advance our understanding of the molecular complexity underlying sarcopenia and functional decline during aging, underscore the value of integrative approaches for studying complex biological processes, and suggest potential molecular targets for future interventions aimed at preserving muscle health and mobility in aging populations.

## Acknowledgements

The Wake Forest Vervet Research Colony is supported by NIH P40 OD010965 with this specific cohort of aging vervets supported by R24 AG073199 (to CAS). Funding for the biopsies and multi-omics data generation was provided by a Wake Forest Claude Pepper Older Americans Independence Center (P30 AG021332) Pilot (to EEQ). Data analysis was supported by R01 AG087957 (to EEQ).

## Statements and Declarations

The authors declare no competing interests.

