## Supplemental Materials for "Integrative Multi-Omics Analysis Reveals Molecular Signatures of Age-Related Decline in Vervet Leg Skeletal Muscle"

GeroScience: Official journal of the American Aging Association

Maitree Patel[1,2*], Eleanor V. Rudasi[1,2*], Brett Frye[3,4], Jacob Negrey[3,5], Laura A. Cox[1,2,3], Ge Li[2], Matthew Jorgensen[3], Thomas C. Register[3], Kylie Kavanagh[1,3,6], Carol A. Shively[3], Ellen E. Quillen[1,2]

1. Section of Molecular Medicine, Department of Internal Medicine, Wake Forest University School of Medicine, Winston Salem, NC
2. Center for Precision Medicine, Wake Forest University School of Medicine, Winston-Salem, NC
3. Section of Comparative Medicine, Department of Pathology, Wake Forest University School of Medicine, Winston Salem, NC
4. Department of Biology & Biochemistry, Emory & Henry University, Emory, VA 24327
5. School of Anthropology, University of Arizona
6. College of Health and Medicine, University of Tasmania, Hobart Australia

*These authors contributed equally to the project.

Corresponding Author: Dr. Ellen E. Quillen; Section of Molecular Medicine, Department of Internal Medicine and Center for Precision Medicine Wake Forest University School of Medicine, Winston-Salem, NC (email)

### Supplementary Materials

#### Individual Omic Feature Analyses Methods

Differential methylation analysis of normalized M-values between normal and declining gait speed (N = 14 per group) was performed using limma [1]. Principle component analysis (PCA) was performed on the cell type proportions obtained from CIBERSORTx, and the first two PCs accounting for 57.03% of the total variance were included as covariates (~ Gait_Status + PC1 + PC2). CpGs were interpreted as hypermethylated if l the difference in M between the groups was greater than 1; and hypomethylated if less than -1 following empirical Bayes moderation which is equivalent to an adjusted p-value < 0.05.

Differential expression (DE) analysis of mRNA and miRNA was performed using DESeq2 [2] with the same covariates as methylation analysis. Differentially expressed mRNA or miRNA were assessed using thresholds of log2 Fold Change (logFC) > 2 for upregulated and logFC < -2 for downregulated transcripts with an adjusted p-value < 0.05. miRNA-mRNA target pairings were evaluated using QIAGEN IPA (QIAGEN Inc., https://digitalinsights.qiagen.com/IPA) to evaluate biological effects of miRNAs. LogFC, p-value, and adjusted p-value columns from miRNA DE results were used as the input without applying any thresholds. Pairings predicted using the Human TargetScan [3] and results with experimental validation were retained. Furthermore, miRNA target prediction was paired with mRNA DE results (logFC) to prioritize miRNA with gene targets present in our data and pairs with opposing expression signals (i.e. upregulation of miRNA and downregulation of its target mRNA and vice versa).

**Individual Omic Feature Analyses Results**

To assess molecular differences using traditional single-omics approaches, DE analysis of mRNAs and miRNAs, and differential methylation analysis of CpG sites were performed between the Normal and Declining groups while adjusting for cell type heterogeneity. mRNA DE analysis using DESeq2 [2] identified 75 DE (absolute logFC > 2) transcripts with nominal p-value < 0.05. After multiple testing correction using the False Discovery Rate (FDR), only *MYL10* was significantly DE (FDR < 0.05, absolute LogFC > 2) between the gait speed groups (Suppl. Fig1a). It is highly expressed in skeletal muscle and has been predicted to enable calcium ion binding and predicted to support contractile force generation. Kong, Wu [4] identifies the gene as a potential indicator for slow-twitch muscles in chickens and therefore with a LogFC ~ -4, its downregulation points to a plausible functional shift in skeletal muscles of vervets with declining gait speed. Two additional genes *CD48* and *CLEC12A* were significant with logFC > 1.7 but both genes have immune-related functions. Since our phenotype under investigation is strongly correlated with age, these two genes might reflect changes associated with chronological age rather than gait speed. Lastly, differential methylation analysis revealed ten significantly differentially expressed CpG sites, out of which only two had a mean difference in $\beta$ > 1, cg20410171 and cg11833660; a third CpG, cg00077017, with a $\Delta\beta$~ 0.93 was also considered significantly differentially methylated.

miRNA DE analysis was also performed using DESeq2 revealing 39 transcripts with unadjusted p-value < 0.05. After FDR correction, two miRNAs were significantly differentially expressed between the groups (Suppl. Fig1b), hsa-miR-100-5p (log2FC ~ -0.55 and adjusted p-value ~ 0.015) and hsa-miR-223-3p (log2FC ~ 1.47 and adjusted p-value ~ 0.007). Evaluating miRNA-mRNA pairings using the criteria described in methods section above, we found that miR-100-5p targets *EPDR1* and *FGFR3*; and miR-223-3p targets *LMO2* and *MEF2C* genes. Despite the confident pairing evidence and discordant logFC results between the pairs; none of the mRNAs reached nominal statistical significance in DE analysis. We also performed Pearson's correlation between miRNA-mRNA raw counts and found no significant correlations.

Despite identifying sparse biological signal from the single-omics analyses, the findings are novel molecular targets for skeletal muscle decline. *MEF2C* is involved in skeletal muscle development [5, 6] and is associated with BMD [7]. Most notably, MEF2C might directly affect gait speed by regulating sarcomeric integrity, power output through fiber-type specification, and overall energy output. EPDR1 and miR-100-5p-FGFR3 pairing jointly highlight bone and musculoskeletal biology. Highly expressed in muscles and as a key potential regulator of bone formation [8], EPDR1 represents common muscle-bone genetic architecture [9]. IPA places miR-100-5p-FGFR3 within the bone mineralization signaling pathway, positioned upstream of calcium reabsorption biological process. While existing literature on FGFR3 focuses on its role in cancers [10-

12] and skeletal dysplasia [13-15], our findings highlight its epigenetic regulation as a potential novel driver of muscle decline in sarcopenia models.

Differential methylation analysis performed using limma [1] revealed 82,424 CpG sites with nominal p-value < 0.05. After FDR correction, ten CpG sites were significantly differentially methylated (FDR < 0.05) between the Normal and Declining groups (Figure 3). Two out of these ten had a mean difference in $\beta > 1$, cg20410171 and cg11833660; a third CpG, cg00077017, with a $\Delta\beta \sim 0.93$ was also considered significantly differentially methylated. All three significant CpGs are hypermethylated in declining animals. Two of the three were able to be annotated with the nearest gene based on probe sequence similarity to the vervet genome. They were both found near genes regulating actin cytoskeletal dynamics. cg11833660 is proximal to the *SSH1* (slingshot homolog 1 protein phosphatase) gene and cg00077017 near the *FLNB* (filamin B) gene. cg20410171 hypermethylation, which is the top signal, was not mapped to a chromosome in the current vervet genome and therefore was not annotated. The Infinium Methylation EPIC Chip Manifest file annotation for its probe sequence was *ZNF578* based on the human genome. *ZNF578* is involved in transcriptional regulation specific to RNA polymerase II. However, a nucleotide blast on the probe sequence against the vervet genome resulted in discordant results as the accession associated with the top hit was linked to multiple zinc finger genes including *ZNF578, ZNF534, and ZNF528*.

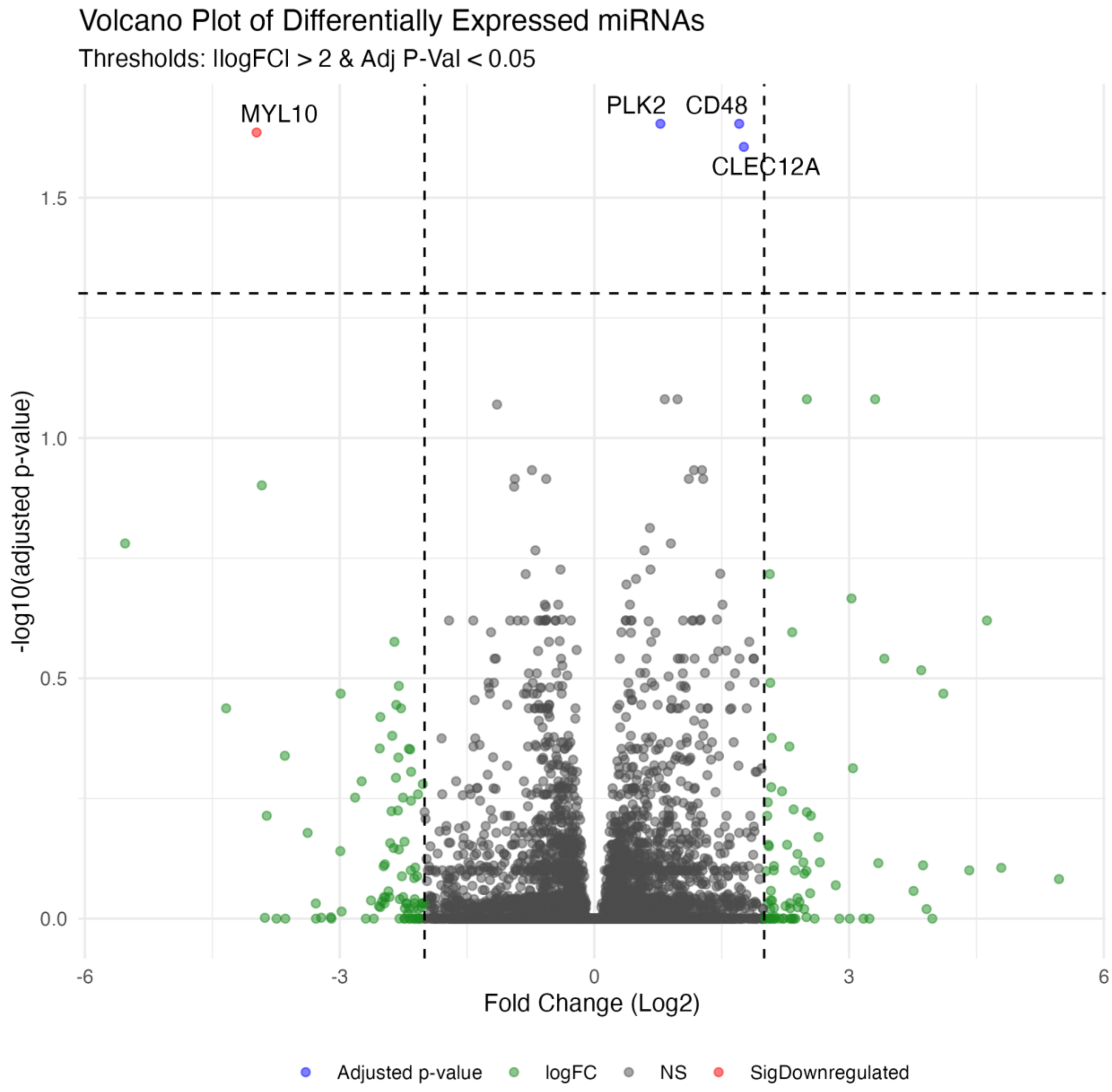


**Supplementary Fig1a** Volcano plot highlighting differentially expressed mRNA between vervets with normal and declining gait status groups, adjusted for cell type proportions using DESeq2. Each point represents a gene, plotted by log2 Fold Change (x-axis) versus statistical significance (-log10 adjusted P-value, y-axis). The horizontal dashed black line indicates the FDR significance threshold (FDR = 0.05), and vertical dashed black lines mark a log2 Fold Change of ±2. The green points represent CpGs with |logFC| > 2 but do not reach FDR significance. The blue points highlight significant miRNAs that are not differentially expressed. Red points show significantly differentially expressed miRNA. The top genes ordered by adjusted p-value are labeled.

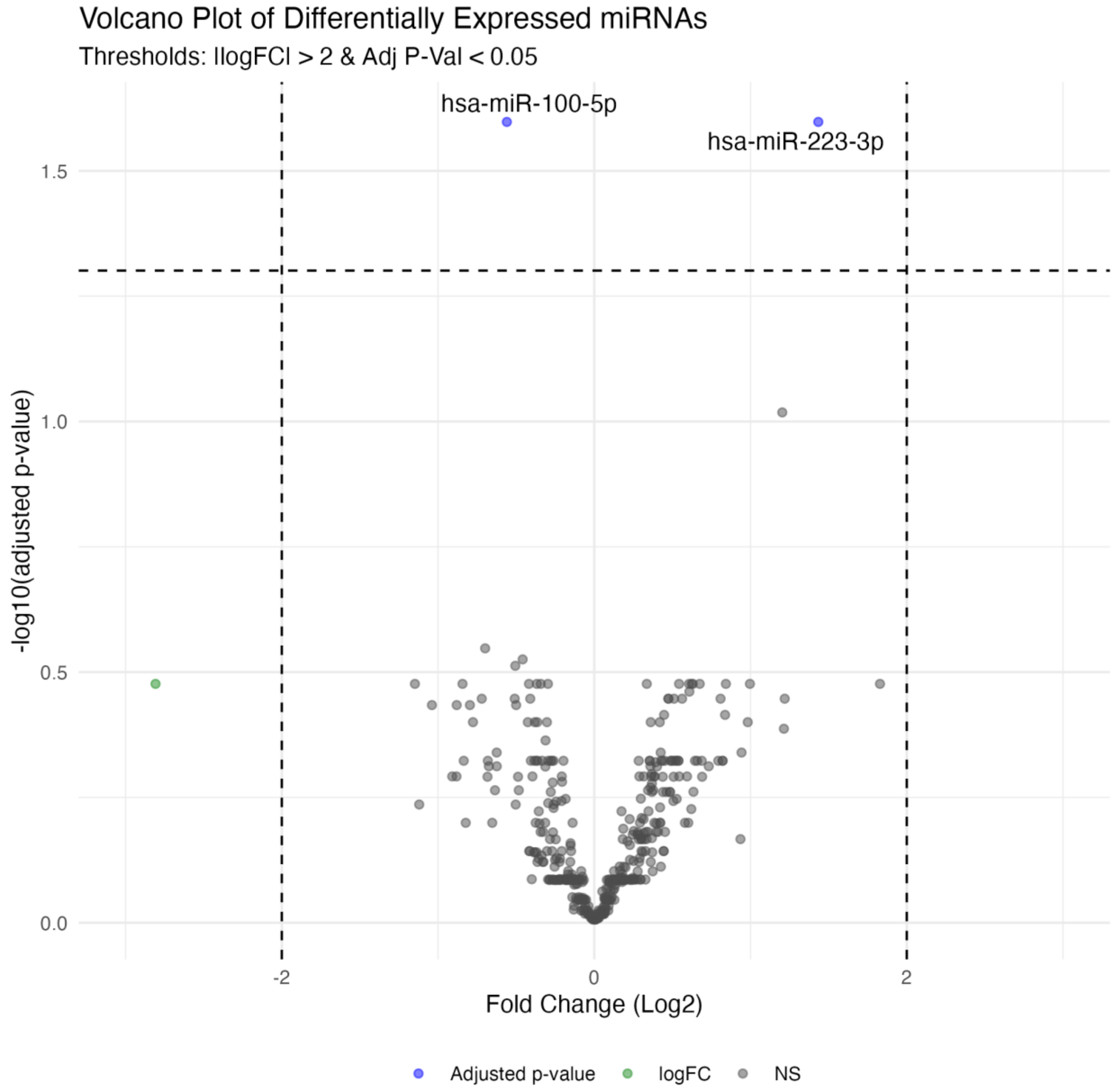


**Supplementary Fig1b** Volcano plot highlights differentially expressed miRNA between vervets normal and declining gait status groups, adjusted for cell type proportions using DESeq2. Each point represents a gene, plotted by log2 Fold Change (x-axis) versus statistical significance (-log10 adjusted P-value, y-axis). The horizontal dashed black line indicates the FDR significance threshold (FDR = 0.05), and vertical dashed black lines mark a log2 Fold Change of ±2. The green points represent CpGs with |logFC| > 2 but do not reach FDR significance. The blue points highlight significant miRNAs that are not differentially expressed. Red points show significantly differentially expressed miRNA. The top miRNAs ordered by adjusted p-value are labeled.

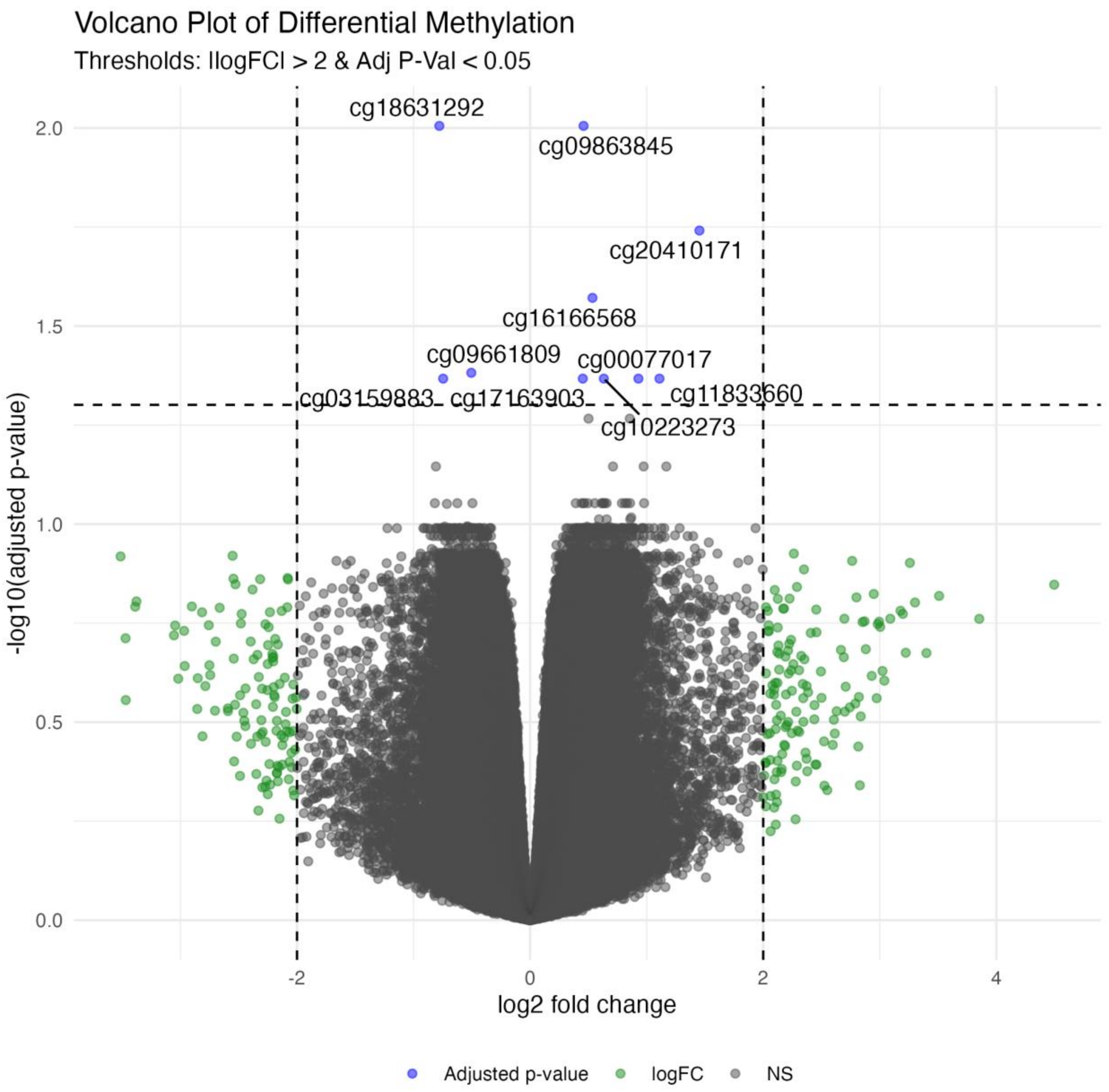


**Supplementary Fig1c** Volcano plot highlighting significant differentially methylated CpG sites between vervets normal and declining gait status groups, adjusted for cell type proportions using DESeq2. Each point represents a gene, plotted by log2 Fold Change (x-axis) versus statistical significance (-log10 adjusted P-value, y-axis). The horizontal dashed black line indicates the FDR significance threshold (FDR = 0.05), and vertical dashed black lines mark a log2 Fold Change of ±2. The green points represent CpGs with |logFC| > 2 but do not reach FDR significance. The blue points highlight significant miRNAs that are not differentially expressed. Red points show significantly differentially expressed miRNA. The top CpG sites ordered by adjusted p-value are labeled.

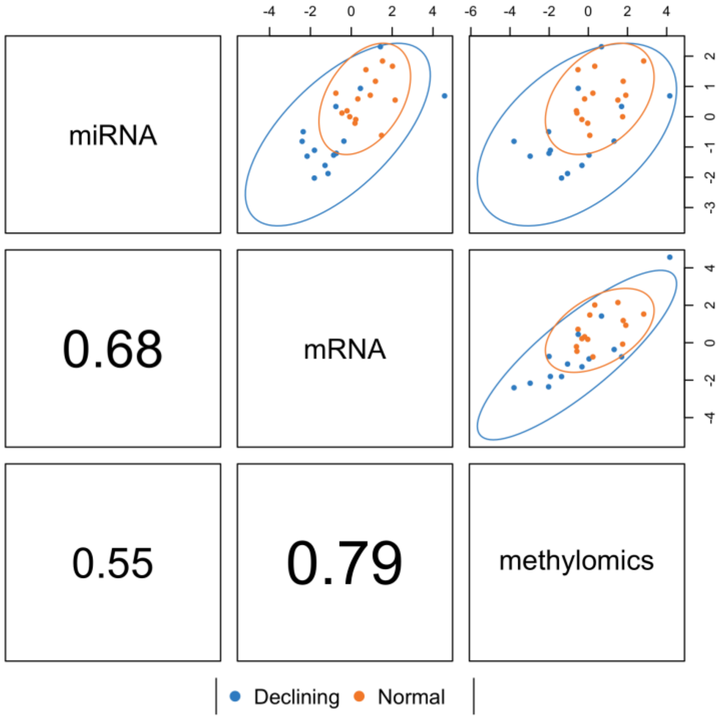


**Supplementary Fig2** Diagnostic plot (plotDiablo from the mixOmics package) visualizing the final DIABLO model integrating mRNA, miRNA, and DNA methylation data. Upper triangle panels: Scatter plots show individual samples projected onto the first two components derived by the model, viewed within each omics block. Samples are colored by gait status group (Declining = blue, Normal = orange), with 95% confidence ellipses indicating group separation achieved based on the integrated components. Lower triangle panels: Pearson correlation coefficients (R values) between the sample scores (variates) on the first component for each pair of omics blocks.

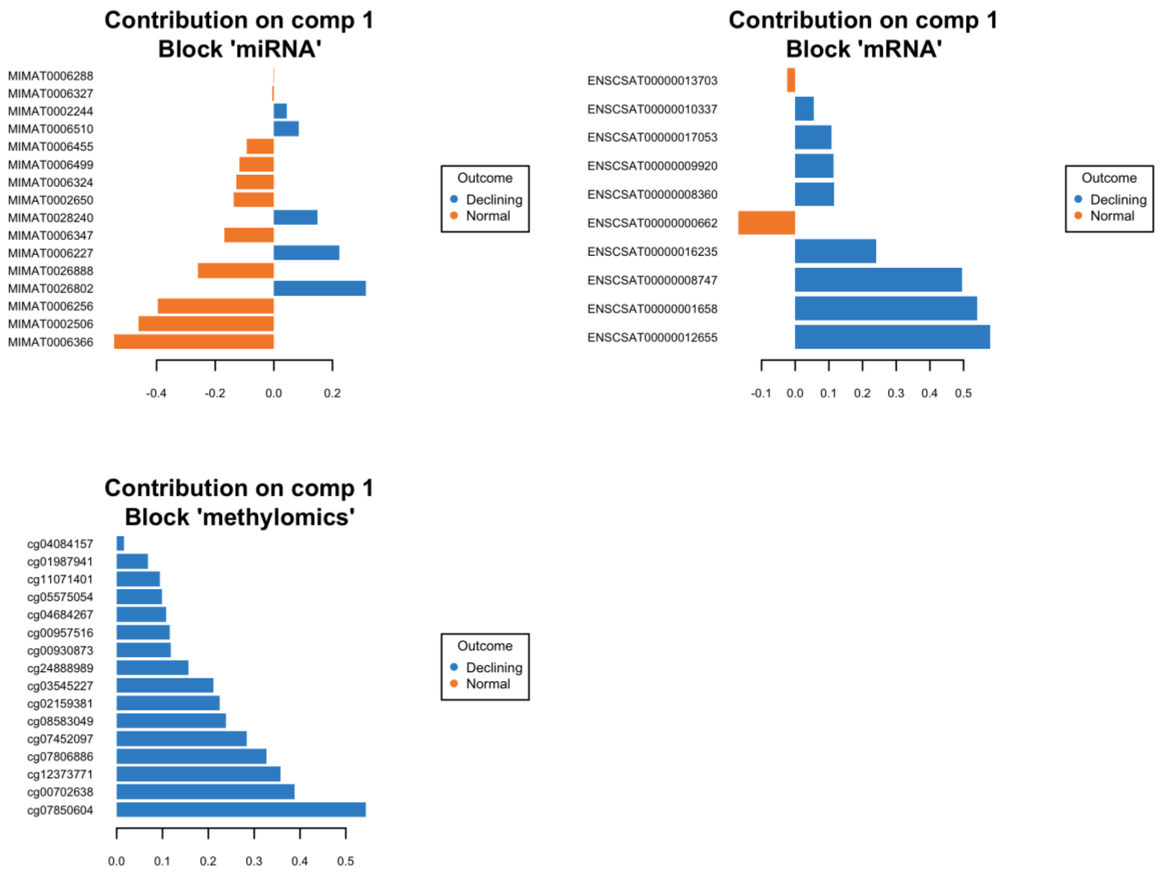


**Supplementary Fig3a** Bar plots illustrating the loading weights of the top features selected by Component 1 of the final DIABLO model, shown separately for each omics block: (A) miRNA, (B) mRNA, and (C) DNA methylation. Features are listed on the y-axis, ordered [confirm ordering, e.g., by absolute loading weight or position on plot]. The loading weight (x-axis) indicates the importance and direction of each feature's contribution to Component 1. The color of the bars indicates the group (Outcome: Blue = higher average level in Declining group; Orange = higher average level in Normal group) in which the feature exhibits higher relative abundance (expression or methylation). These plots identify the specific molecular features across the different omics layers that most strongly contribute to the primary latent component distinguishing the gait speed groups.

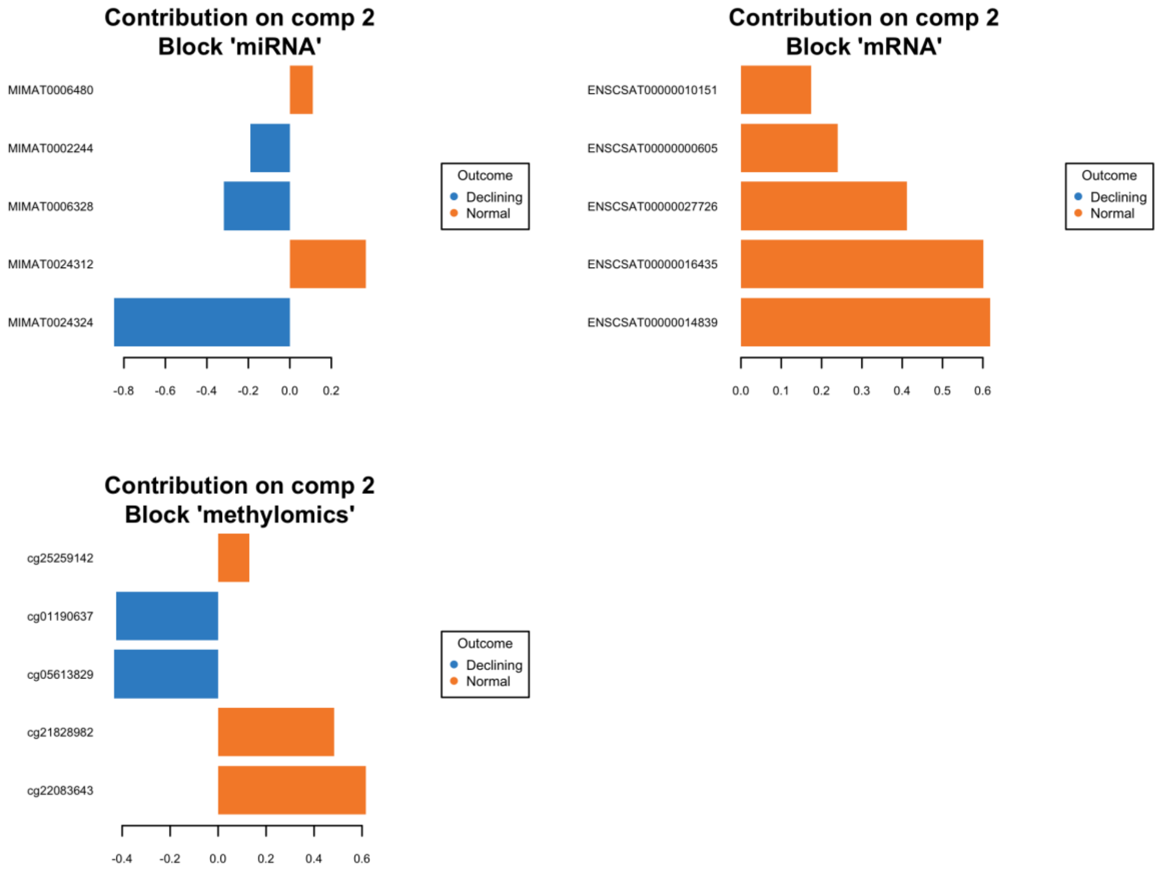


**Supplementary Fig3b** Bar plots illustrating the loading weights of the top features selected by Component 2 of the final DIABLO model, shown separately for each omics block: (A) miRNA, (B) mRNA, and (C) DNA methylation. Features are listed on the y-axis, ordered [confirm ordering, e.g., by absolute loading weight or position on plot]. The loading weight (x-axis) indicates the importance and direction of each feature's contribution to Component 2. The color of the bars indicates the group (Outcome: Blue = higher average level in Declining group; Orange = higher average level in Normal group) in which the feature exhibits higher relative abundance (expression or methylation). These plots identify the specific molecular features contributing most significantly to the second latent component, which captures additional variation discriminating between the gait speed groups.

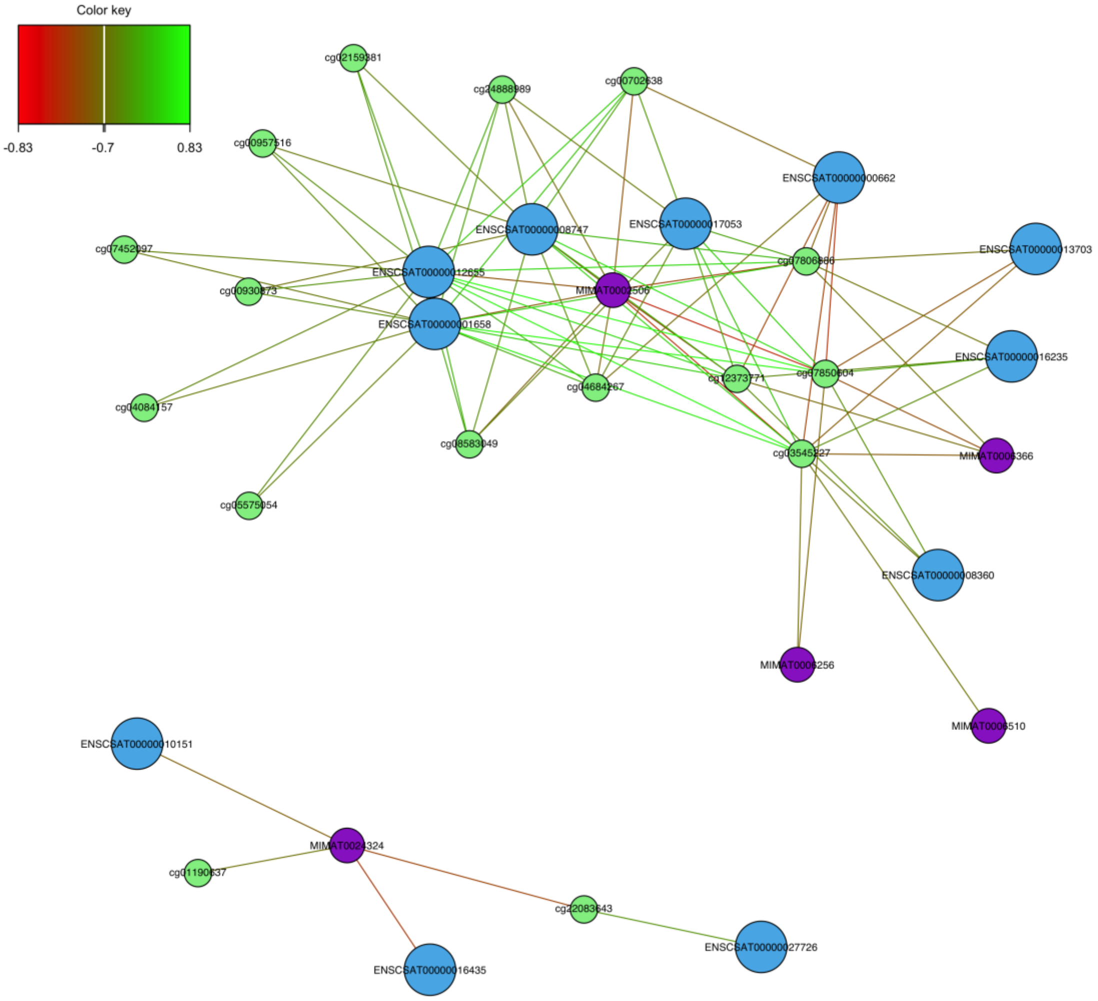


**Supplemental Fig4** Network visualization of the relationships between key features selected by the final two-component DIABLO model from the mRNA (light blue nodes), miRNA (purple nodes), and DNA methylation (light green nodes) datasets. Nodes represent individual features (labeled by ID). The color intensity within each node reflects its correlation with Component 1 (Color key: Red = negative correlation, Green = positive correlation, range -0.9 to 0.9). Edges (lines) connect pairs of features whose absolute Pearson correlation exceeds a threshold of 0.7. Edge color indicates the sign of the correlation between connected features (Green = positive, Red = negative). The network structure reveals two distinct clusters, primarily representing features contributing to Component 1 (larger, top cluster) and Component 2 (smaller, bottom cluster), highlighting the interconnected molecular signature associated with gait speed status.
